# Reactivation of latent HIV-1 *in vitro* using an alcoholic extract from *Euphorbia umbellata* (Euphorbiaceae) latex

**DOI:** 10.1101/293662

**Authors:** Ana Luiza Chaves Valadão, Paula Pezzuto, Viviane A. Oliveira Silva, Barbara Simonson Gonçalves, Átila Duque Rossi, Rodrigo Delvecchio da Cunha, Antonio Carlos Siani, João Batista de Freitas Tostes, Marcelo Trovó, Paulo Damasco, Gabriel Gonçalves, Rui Manuel Reis, Renato Santana de Aguiar, Alves de Melo Cleonice Bento, Amilcar Tanuri

## Abstract

*Euphorbia umbellata (E. umbellata)* belongs to *Euphorbiaceae* family, popularly known as Janauba, and contains in its latex a combination of phorbol esters with biological activities described to different cellular protein kinase C (PKC) isoforms. Here, we identified deoxi-phorbol esters present in *E. umbellata* latex alcoholic extract able to increase HIV transcription and reactivate HIV from latency models. This activity was mediated by NF-kB activation followed by nuclear translocation and binding to HIV LTR promoter. In addition, *E. umbellate* latex extract induced the production of pro inflammatory cytokines together with IL21 in *in vitro* human PBMC cultures. Our latex extract activates latent HIV in human PBMCs isolated from HIV positive patients as well as latent SIV in non-human primate primary CD4^+^ T lymphocytes. These results strongly indicate that the phorbol esters present in *E. umbellata* latex are promising candidate compounds for future clinical trials for *shock and kill* therapy to promote HIV cure and eradication.

## INTRODUCTION

The Acquired Immunodeficiency Syndrome (AIDS) is caused by the Human Immunodeficiency Virus (HIV), which was identified in 1983 (**1,2**) from patients with AIDS. In those 34 years of pandemic, more than 35 million people have died from AIDS-related illnesses since the start of the epidemic until 2016 (**3**).

The pandemic of HIV infection is an important socioeconomic burden and is considered one of the largest documented epidemics in history. HIV pathogenesis involves infection and replication in CD4^+^ T lymphocytes, macrophages and dendritic cells. Replication and destruction of CD4^+^ T cells, which are key effector cells in the host immune response, leads to a clinical outcome of immunosuppression known as AIDS (**4**).

The current strategy used in the treatment of HIV infection is named Antiretroviral Therapy (ART), which is based on a combination of antiretroviral drugs, which target different stages of HIV replicative cycle. The primary goal of ART is to block viral replication and slow down the progression of immunodeficiency, restoring immunity and improving the quality of life of the infected patient (**5**). However, adverse effects have been reported as a consequence of the antiretroviral therapy, in conjunction with immunological, neurological and metabolic co-morbidities associated either with HIV infection or drugs interaction (**6**). These facts triggered the search for a functional or sterilizing cure for HIV infection.

Even when an undetectable viral load is achieved with effective antiretroviral therapy, it is still possible to detect latent viral reservoirs where HIV proviral DNA is integrated in resting memory CD4+ T cells (**7**). These cells containing latent viral genomes become viral reservoirs inaccessible to ART that can only act on actively replicating viruses. Once ART is stopped, viral load increases rapidly and systemic infection reestablishes (**8**). Therefore, the complete eradication of HIV from an infected individual is one of the most challenging areas of HIV research today. The way to eliminate HIV from patient is to activate the virus in the presence of ART and or activate the immune response to destroy viral reservoir or make patient´s immune response to keep viral replication at the bay.

New latent virus reactivating compounds, also called latency reversing agents (LRA), when used conjointly with ART, are capable to eliminate transcriptionally inactive viruses, ultimately resulting in the eradication of HIV infection. Histone deacetylase (HDAC) inhibitors are promising classes of latency-reversing agents (LRAs) that are undergoing extensive testing in *in vitro* models and in initial pilot clinical trials to reactivate latent HIV-1 infection. HDAC inhibitors were developed as anticancer drugs as HDACs play important role in epigenetic and non-epigenetic transcriptional regulation, inducing apoptosis and cell cycle arrest (**9**). In the context of HIV-1 reactivation, HDAC inhibitors induce transcription at the HIV-1 long terminal repeat (LTR) (**10–13**). This strategy is called “shock and kill” and it postulates that memory CD4+ T lymphocytes treated with LRA along with ART could purge totally or partially the viral reservoir while blocking replication of emergent latent virus, leading to a HIV eradication or at least a functional cure (**14**). Another promising class of molecules with therapeutic potential has been reported as possible candidates for elimination of viral reservoirs are the Protein Kinase C (PKC) agonists (**7**). This class includes: Prostratin (**15**), Bryostatin (**16**), Ingenol (**17**) and Phorbol 12-Myristate 13-Acetate (PMA) (**18**). These compounds can reactivate latent HIV in lymphoid and myeloid cells. Although several transcriptional regulatory mechanisms for HIV latency have been described in the context of epigenetic silencing and transcription repression, recent work suggests that reactivation of latent HIV is positively correlated with NF-κB activation, demonstrating the importance of this factor in latency reactivation (**19, 20**).

Many natural compounds have been investigated for their antiviral and latency-reversing properties, being indicated as candidates for clinical trials. Several of these compounds are derived from secondary metabolics of plants, such as terpenoids (**21**), polyphenols (**22**), alkaloids (**23**) and phorbol esters (**24**). *Euphorbiaceae* family is a source of potent phorbol esters, which specimens are found worldwide and havebeen used for the treatment of several diseases in the popular medicinal use (**25–29**). *Euphorbia umbellate* is popularly known in Brazil as Janauba or *leiterinha* and is used, in traditional medicine, as a latex preparation (diluted in water) for the treatment of gastric disorders, such as peptic ulcer and gastritis, as well as neoplastic diseases, allergy, diabetes, influenza, leprosy, among other diseases (**30**). Several papers have reported the isolation of lectins, phorbols, terpenes, flavonoids and other phenolic compounds from *E. umbellata* latex (**31–33**). This plant contains phorbol esters in its latex similar to PMA isolated from the seeds of *Croton tiglium,* another representative of the *Euphorbiaceae* family. These compounds are efficient in inhibiting the cytopathic effect generated by HIV infection on lymphocyte cells, as well as activating protein kinase C (PKC) that in turn activates NF-κB, promoting its nuclear translocation and promoting HIVLTR transcriptional reactivation reverting viral latency (**34**).

In this work we evaluated the ability of an alcoholic extract obtained from Janauba latex (JALEx) to reactivate latent HIV and investigated the molecular mechanisms involved in this activity. The anti-latency property of JALEx was demonstrated either in *ex vivo* HIV latent derived human PBMC cells or in CD4+ T Lymphocytes isolated from SIV infected non-human-primates. All these properties makes JALEx a promising natural medicine to be used in HIV cure strategies.

## MATERIAL AND METHODS

### Latex isolation from *Euphorbia umbellata*

*Euphorbia umbellata* (Pax) Bruyns, the current accepted name for *Synadenium grantii* Hook.f., was collected and identified with the respective voucher specimen kept at the RFA Herbarium under the number Trovó, M.L.O 764. The crude latex was collected directly from incision mechanically made in the trunk of cultivated specimens at Barretos county, São Paulo state, Brazil. The latex was straightly diluted in 80% ethanol (1:10 v/v) to produce a white rubber-like precipitate that was filtered off by gravity in filter papers. The filtrate was dried in a rotary evaporator to afford a fine white powder, which was diluted in DMSO to a final concentration of 20 mg/mL. This extracted material was labeled as JALEx (Janaúba Alcoholic Latex Extract). JALEx was chromatographed in a silica gel column using a solvent gradient (0-5%) of ethyl ether in chloroform to enrich the phorbol-containing fractions and favor their detection and characterization.

### Chemical profile of JALEx

JALEx was analyzed by gas chromatography coupled to mass spectrometry (GC-MS) by using an Agilent 6890N chromatograph (Palo Alto, CA) equipped with a capillary column DB-H17HT (30 m × 0.25 mm × 0.15 μm film thickness). Helium was the carrier gas at 1.0 mL/min and inlet pressure of 3.14 psi, 10:1 split/splitless ratio. The oven temperature programmed from 150°C to 340°C at 4°C/min and then held for 17 minutes. The injection volume was 1 μL of a 2 mg/mL chloroform solution. Data were processed using MSD Productivity Chem Station Software operating with ion source at 250°C and electron impact ionization at 70 eV. Individual components were characterized by their fragmentation patterns in the mass spectrum.

### Viruses and cells

Analysis of HIV reactivation *in vitro* in latent infected cells after treatment with JALEx. Human HIV latent lymphocyte cell models (J-Lat 8.4 and 10.6) were used (**35**). These cells are derived from Jurkat cells containing copies of HIV genome in fusion with GFP reporter gene integrated into the nuclear. These cells have different levels of reactivation capacity, depending on the site of HIV genome integration. The J-Lat 10.6 clone is a more responsive one showing reactivation levels higher than those observed by the J-Lat 8.4 clone. Both cells are cultured in RPMI medium supplemented with 10% fetal bovine serum (FBS) and 1% penicillin / streptomycin antibiotics. J-Lat cells at density of 10^5^ cells per well of 24-well plates were treated with different concentrations of JALEx dissolved in DMSO for 24 hours. Positive controls for reactivation of HIV latent virus were ingenol B (0.32 μM) and PMA (1 μM). Cells were analyzed by flow cytometric (BD Accuri C6) to determine the level of GFP expression, in order to evaluate HIV transcription. A total of 10,000 gated live cells were collected and data represent the percentage of GFP-expressing cells in total gated events. All the experiments were performed in three independent replicates.

### Cell viability assay

Evaluation of the cytotoxicity of the compounds was performed by incubating J-Lat cells with the serial dilutions of JALEx compounds previously tested in the reactivation assays. Cell viability assessment was performed in 10^4^ cells plated in 96-well plates, in three independent experiments, each in sextuplicates. After 5 days of culture, vital resazurin dye (CellTiter Blue, Promega) was added to the cells and absorbance levels were evaluated by spectrophotometer at 595 nm.

### Analysis of the activation of PKCs

To assess the mechanism of action of HIV latency reactivation by Janauba compounds, 10^5^ J-Lat cells in 24-well plates were treated with different specific inhibitors of PKC**α**, PKC**δ** and PKC**γ** isoforms Go6976, Go6983, Ro-31-8220, respectively, for 24 hours (**36–38**). After this time, cells were treated with the positive control of PKC activation (1 μM PMA) and two concentrations of JALEx previously determined in the reactivation and cytotoxicity assays. Untreated cells were referred as MOCK. The following concentrations of the alcoholic extract were tested: 0.01 μg/mL and 0.001 μg/mL and the percentage of GFP positive cells was assessed by flow cytometry analysis.

In addition, PKC activation was evaluated by immunofluorescence to detect cytosolic location of different PKC isoforms. For this, 10^4^ HeLa cells were plated in black 96-well glass bottom plates. These cells were transfected with three different plasmids encoding three PKC isoforms fused to GFP (PKC**α**-GFP, PKC**δ**-GFP and PKC**γ**-GFP) for 24 hours and then treated with 1 μg/mL and 0.1 μg/mL of the JALEx, for 10 minutes or 24 hours. Positive control was performed using PMA (1 μM). Cells were washed with PBS and DAPI stained. Cellular localization of PKC-GFP was evaluated by High Content Screening confocal microscope (Molecular Devices, Inc). The redistribution of PKC isoforms was quantified in ImageJ software.

Phosphorylation of PKC isoforms was also investigated. For this, Jurkat cells were treated with 0.01 μg/mL and 0.001 μg/mL of JALEx and with 1 μM PMA as a positive control for 10 and 30 minutes, 1, 6 and 24 hours. Cellular extractes were prepared in lysis buffer (50 mM Tris pH 7.6, 150 mM NaCl, 5 mM EDTA, 1 mM Na_3_VO_4_, 10 mM NaF, 10 mM NaPyrophosphate, 1% NP-40 supplemented with protease inhibitors cocktail) for 1 hour. Lysates were subjected to 10% SDS-PAGE gels and after transferred to nitrocellulose membranes the following antibodies were used for immunoblotting: anti-PKC phosphorylated PKC (pan, **δ**, θ) (Cell Signaling; 1:500) and **β**-tubulin (Cell Signaling; 1:2000). Proteins quantification was evaluated using band densitometry analysis, performed with the Image J software (version 1.41; National Institutes of Health).

### NF-κB activation assays

In order to verify whether JALEx induces the activation and downstream nuclear internalization of NF-κB, HeLa cells were plated at the density of 2×10^4^ cells per well in 96-well plates. Cells were treated with 1 μg/mL and 0.1 μg/mL of JALEx for 6 or 24 hours. PMA (1 μM) was used as positive control. The NF-κB subunit p65 was visualized by immunofluorescence (anti-p65, 1:300, Santa Cruz Biotechnologies), and nuclear (DAPI) translocation was evaluated by fluorescence confocal microscopy (Molecular Devices, Inc).

In order to evaluate the role of NF-κB in reactivation of latent HIV, Jurkat cells (10^6^ cells) were transiently transfected with 8 μg of pBlue30LTR-Luc Wild Type (NFkB WT) or p-LTR-MUT (NFkB MUT) using the Neon Transfection System kit (Thermo Fisher Scientific) according to the manufacturer’s recommendations. The latter plasmid was generated by site-directed mutagenesis by altering the two NF-κB binding sites, rendering that plasmid nonfunctional for NF-κB response. Both plasmids were provided by Dr. Renato S. de Aguiar (UFRJ) and those constructs have the luciferase sequence under the control of the HIV viral promoter. In the case of plasmid carrying NF-kB mutate sequence, the second guanine at both NF-κB binding sites into LTR promoter was changed to a cytosine, resulting in the plasmid designed here as *pBlue3’LTR NF-κB MUT-Luc* (NF-κB BS^−^). This plasmid carries a LTR sequence lacking functional NF-B binding sites. After transfections, cells were equally divided into two groups: treated or untreated with 1μg/mL of JALEx, for 24 hours. After this interval, cells were lysed and luciferase activity was assessed in a luminometer (Glomax, Promega).

### Analysis of surface markers expression and cytokines production in primary cells treated with JALEx

In order to expand the results obtained in HIV latency model cells, we performed assays with JALEx in peripheral blood mononuclear cells (PBMCs) from 4 healthy donors. After separation of buffy coat by Ficoll centrifugation, CD4^+^ T cells were isolated using the Dynabeads CD4 Positive Isolation Kit (Thermo Fisher) kit to obtain 10^7^ PBMCs, according to the manufacturer’s recommendations. TCD4^+^ cells were treated with 1 μM PMA, 10 μg/mL (10^−3^) or 1 μg/mL (10^−4^) of JALEx, in addition to MOCK (untreated cells), for 24 hours. Expression of surface markers such as CD4, CCR5 and CXCR4, as well as the expression of activation markers CD25, CD38, HLA-DR and CD69 were evaluated by specific antibody labeling and flow cytometric analysis (BD Accuri C6). Comparatively, this assay was also performed on MT4 lymphocyte cells line at the same conditions.

Cell supernatants from the TDC4^+^ cells were collected for analysis of the cytokine production stimulated by JALEx treatment. A colorimetric multiplex immunoassay Bio-Plex Pro Human Cytokine 17-plex Assay (Bio-Rad) containing fluorescent microspheres conjugated to different monoclonal antibodies specific for each target cytokine was chosen for this analysis. The following cytokines were measured: IL-1**β**, IL-2, IL-4, IL-5, IL-6, IL-7, IL-8, IL-10, IL- IL-17, G-CSF, GM-CSF, MCP-1 / MCAF, MIP-1**β**, IFN**γ** and TNF-**α**.

Quantitative viral outgrowth assay. To compare the ability of the compounds to stimulate virus replication in latently infected cells we used a variation of the quantitative viral outgrowth assay (QVOA) described by Laird et al. (**39**). Cells for the QVOA were isolated from viremic SIVmac239-infected cynomolgus macques at the Wisconsin National Primate Research Center (WNPRC). PBMC isolated by density gradient centrifugation were enriched for CD4^+^ T cells using the CD4 T cell enrichment kit for non-human primates (Miltenyi Biotec, San Diego, CA). After enrichment, cells were plated at 1×10^6^ cells per well. These cells were co-cultured with “stimulator” CEMx174 cells that were irradiated with 10,500 rads prior to co-culture. The ratio of stimulator cells to CD4^+^ T cells was 2:1. Purified CD4^+^ T cells were stimulated with JALEx, concanavalin A, ingenol, or no mitogen, overnight. After stimulation, cells were washed and re-plated in serial dilutions, in triplicate. At this point, additional non-irradiated CEMx174 cells were added to each well to serve as targets for induced virus to infect. The target cells were added at a 2:1 ratio to the CD4^+^ T cells. Co-cultures were maintained for 1 week. On the last day, supernatant was collected and RNA was isolated to test for the presence of SIV. RNA was isolated using the Viral Total Nucleic Acid kit for the Maxwell 16 MDx instrument (Promega, Madison, WI). Viral RNA was quantified using a qRT-PCR assay targeting the *gag* gene as described previously (**40**) except that analysis was performed using the Lightcycler 480.

### Effect of JALEx in T cell activation from HIV positive patients

In order to analyze the ability of JALEx in regulating different T cell phenotypes in the context of HIV-1 infection, peripheral blood of 15 adults ART-treated patients (8 women and 9 men), with a mean plasma viral load of 953 ± 1.924 copies of RNA/mL and CD4^+^ (845 ± 345/mm^3^) and CD8^+^ (845 ± 345/mm^3^) higher than 350 cells/mm^3^, were collected and PBMC were obtained from Ficoll-Paque gradient. Viable PBMC were quantified by trypan blue exclusion and suspended in RPMI-1640 medium supplemented with 2 μM of L-glutamine (GIBCO, Carlsbad, CA, USA), 10% of fetal calf serum, 20 U/mL of penicillin, 20 μg/mL of streptomycin and 20 mM of HEPES buffer. Approximately 1 × 10^6^ PBMC/mL were cultured for 24 hours in 24-well plates with 1mL of complete medium in the presence or absence of different concentration of JALEx (1, 0.1 and 0.01 μg/mL). Phorbol-12-myristate-13-acetate (PMA 20 ng/mL) plus ionomycin (600 ng/mL) were used as positive control. To optimize intracellular cytokine staining, brefeldin A (10 μg/mL) was added to the cultures in the last 4 hours of incubation time. For the fluorescence labeling, mouse anti-human fluorescent monoclonal antibodies (mAbs) directed against CD4-FITC, CD8-FITC were purchased from BD Bioscience (San Diego, CA, USA). Briefly, anti-CD4 and anti-CD8 mAbs were added to the PBMC (2 × 10^5^/tube) and incubated for 30 min at room temperature in the dark. The cells were washed with PBS and then permeabilized with Cytofix/Cytoperm (BD Pharmingen, San Diego, CA) at 4 °C for 20 min. After washing, the antibodies for intracellular staining (anti- IL-17-PECy7, anti-IFN-**γ**-PE-Cy5, anti-IL-10-PE-Cy7, anti-IL-21-PE) or the corresponding anti-IgG1 isotype control, were added in various combinations and incubated for 30 min at 4°C. Cells were analyzed in the Attune^®^ (Thermo Fischer, Co) and FlowJo software. Isotype control antibodies and single-stained samples were periodically used to check the settings and gates on the flow cytometer. After acquisition of 100,000 to 200,000 events, lymphocytes were gated based on forward and side scatter properties after the exclusion of dead cells, by using propidium iodide and doublets. The analysis of CD4^+^ and CD8^+^ T cells viability was determined using 7-aminoactinomycin D (7-AAD) (R&D systems). Further, the gated cells were negative for CD14 marker. The CD4^+^ T lymphocyte count (CD4) was performed on a flow cytometer (FACS Count, Becton Dickson^®^). Plasma HIV RNA quantification was performed using the RT-PCR with Abbott M2000 Sq HIV VL technology.

### Statistical analysis

The statistical analysis was performed using Prism 5.0 software (GraphPad Software). Statistical evaluations were performed by Kruskal-Wallis Test, Mann-Whitney U test and Two-way ANOVA. The analysis are indicated in the figures.

## RESULTS

### JALEx isolation from *Euphorbia umbellate and chemical analysis*

The JALEx soluble in water and ethanol was characterized mainly by a mixture of triterpenes and phorbol ester diterpenes. The GC profile and MS data indicated that triterpenes was predominant throughout the whole extract, with two main constituents standing out, represented as T1 and T2 in figure 1A. The triterpenes profile is in accordance with data from the literature that report the isolation of euphol, lanosterol, friedelin and 3**β**-friedelinol in the latex, leaves and stem of *E. umbellata* (**41**), whilst studies with the latex have led to the characterization of euphol and the steroid citrostadienol (**42**). A phorbol esters-rich fraction was obtained by chromatographing the crude JALEx in a silica gel open column using mixtures of chloroform-ether as eluent (Ph1 and Ph2 - Fig. 1). The class of phorbol ester has been represented by the 12-O-tigloyl-4-deoxyphorbol-13-isobutirate (**43**), 12-deoxyphorbol-13-(2-methylpropionate), phorbol 12,13,20-triacetate (**44**), 4-deoxyphorbol-12,13-ditiglate and 3,4,12,13-tetraacetylphorbol-20-phenylacetate (**45**) distributed in several plant tissues (Fig. S1).

**Figure 1.**
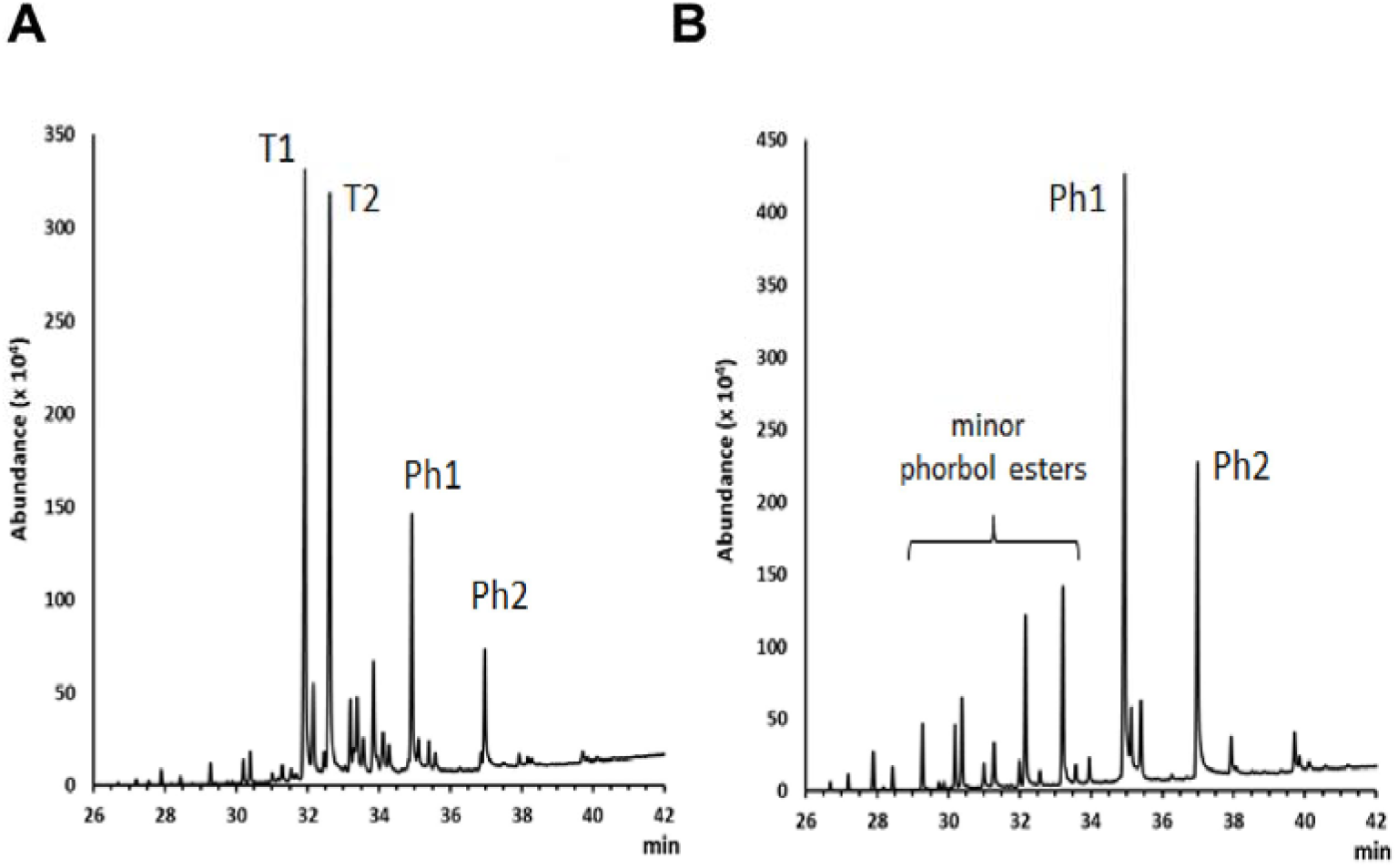
Isolation and chemical structures of JALEx. Gas chromatograms of (A) whole extract JALEx and (B) Phorbol esters-enriched JALEx fractions. T1 and T2 = major triterpenes constituents; Ph1 and Ph2 = major phorbol esters constituents. The chemical classes were indicated by the characteristic mass fragmentation in the GC-MS analysis and data from literature.

### Reactivation of latent HIV by JALEx and its cytotoxicity effects

In this work, we used the alcoholic extracts from JALEx resuspended in DMSO. The reactivation of latent HIV was evaluated using the lymphocytic lineage J-Lat cells (clones 8.4 and 10.6) harboring the HIV integrated genome. The cells were exposed to increasing concentrations of JALEx, ranging from 10 to 10^−7^ μg/mL. The reactivation of latent HIV was evaluated by GFP reporter gene expression cloned in fusion with the virus integrated genome. In parallel, cytotoxicity assay was performed to evaluate whether the concentrations of JALEx would be toxic to these cells. JALEx treatment at concentrations of 10 to 0.01 μg/mL showed HIV activation percentage similar to positive controls (PMA 1 μM and ING-B 0.32 μM), dropping to less than 5% reactivation at dilutions of 0.001 to 0.0000001 μg/mL in the J-Lat 8.4 clone (Fig. 2A). Our results showed cytotoxic effects of JALEx at 10 μg/mL, while the other concentrations were well tolerated, showing a good cell viability (Fig. 2B). As expected, HIV reactivation by JALEx was more expressive in J-Lat cells clone 10.6, reaching values of 75% compared to 12% reactivation in 8.4 cells (Fig. 2C). In relation to the cytotoxicity, JALEx at 10 μg/mL was also toxic for J-Lat 10.6, whereas the other concentrations presented cells viability between 80% and 75% (Fig. 2D). These results demonstrated the ability of JALEx to stimulate HIV transcription and reactivate virus from latency models.

**Figure 2:**
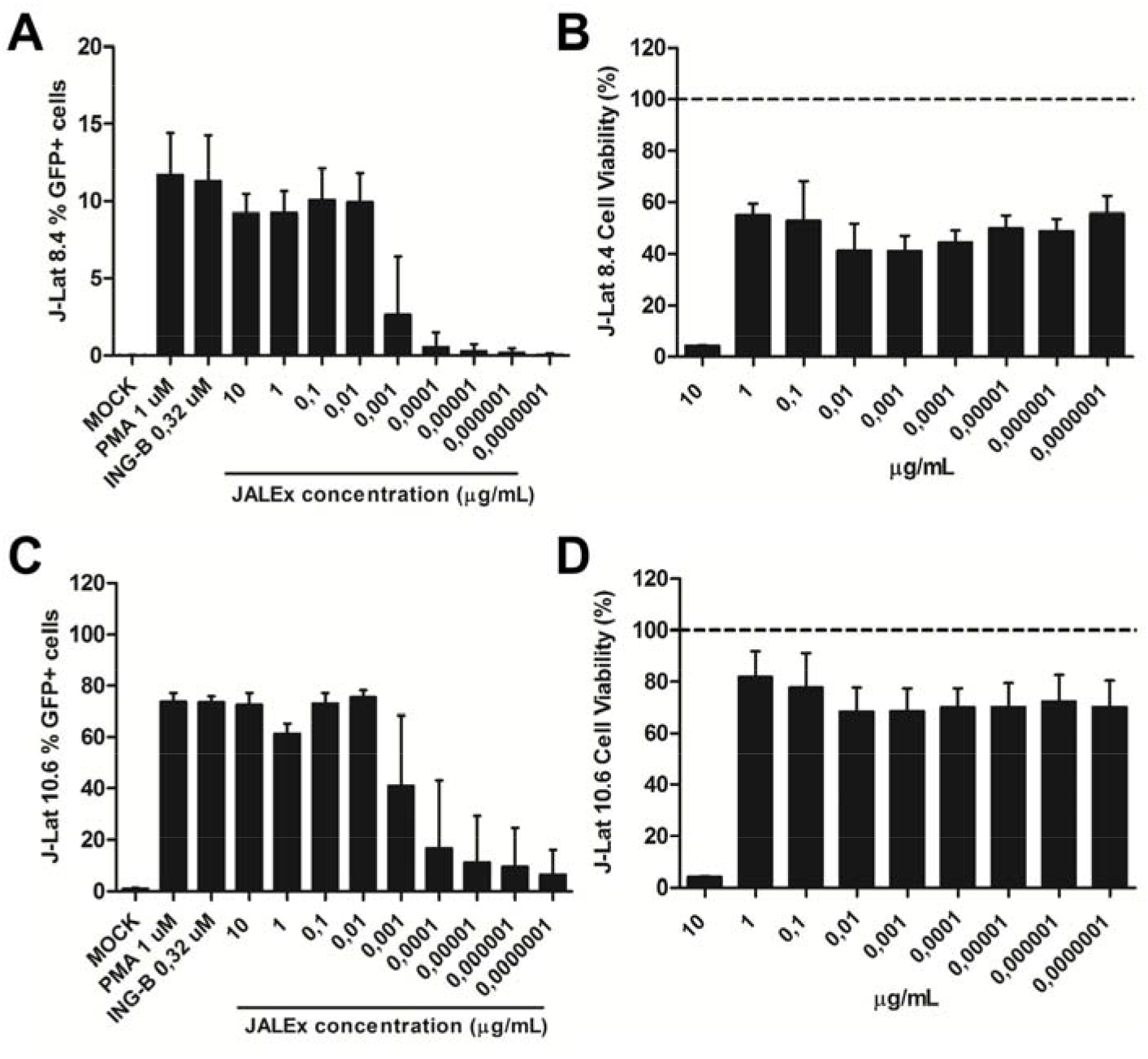
JALEx reactivates virus transcription in HIV-1 latent models. J-Lat 8.4 (A) and 10.6 (C) cells were exposed to increasing concentrations of JALEx and HIV-1 activation was assessed 24 hours post-treatment by GFP positive cells screening by flow cytometry. Mock stands for not treated. PMA (1 μM) and INGB (0,32 μM) were used as positive induction controls. The cytotoxicity of JALEx was evaluated using CellTiter Blue vital stain 5 days after treatment in J-Lat 8.4 (B) and 10.6 cells (D). GFP positivity and cytotoxicity were expressed as percentage of cells normalized by mock non-treated cells (n=3). The dashed lines in Figure 2 B and D represents 100% of viability measures in MOCK. The bars in the figure represents 1 SD around the mean.

### JALEx reactivation of latent HIV is mediated by PKC

In order to evaluate whether JALEx activation of HIV transcription is mediated by PKC pathway J-Lat 8.4 and 10.6 cells were treated for 24 hours with different PKC inhibitors (Gö6666, Gö6983 and Rö31-8220) to prevent JALEx stimulation. After this period, the cells were incubated with non cytotoxic concentrations of JALEx (0.01 and 0.001 μg/mL). PMA is a phorbol ester that activates different isoforms of PKCs and was used as positive control (1 μM). HIV reactivation was evaluated by flow cytometry for quantification of GFP positive cells. The pre-exposition of both clones of J-Lat cells (8.4 and 10.6) to the PKC inhibitors reverse the activation promoted by the PKC agonist PMA compared with PMA only treated cells (Fig. 3). The same results were found when J-Lat clone 8.4 was pre-incubated with inhibitors Gö6976, Gö6983 and Rö31-8220 before JALEx treatment either at concentration 0.01 or 0.001 μg/mL. This result suggests that JALEx is a pan activator of different isoforms of PKCs (PKC**α**, PKC**δ** and PKC**γ**) (Fig. 3 - left panel). The same profile was found with the J-Lat clone 10.6 indicating that PKC inhibitors block the reactivation of latent HIV by JALEx treatment. Rö31-8220 was the only inhibitor that could not totally prevent JALEx stimulation reaching values around 20% of activation compared with 60% of non-treated cells (Fig. 3 - right panel). All together, these results suggests that JALEx is a broad range PKC agonist, since the treatment with PKC inhibitors Gö 6976 (specific for PKC-**α** and **β**), Gö 6983 (specific for PKC-**α**, PKC-**β**, PKC-**γ**, PKC-**δ** and PKC-**ζ**) and Rö-31-8220 (specific for PKC-**α**, PKC-**β**, PKC-**ε** and PKC-**γ**) decreased the levels of HIV reactivation compared with non treated cells.

**Figure 3:**
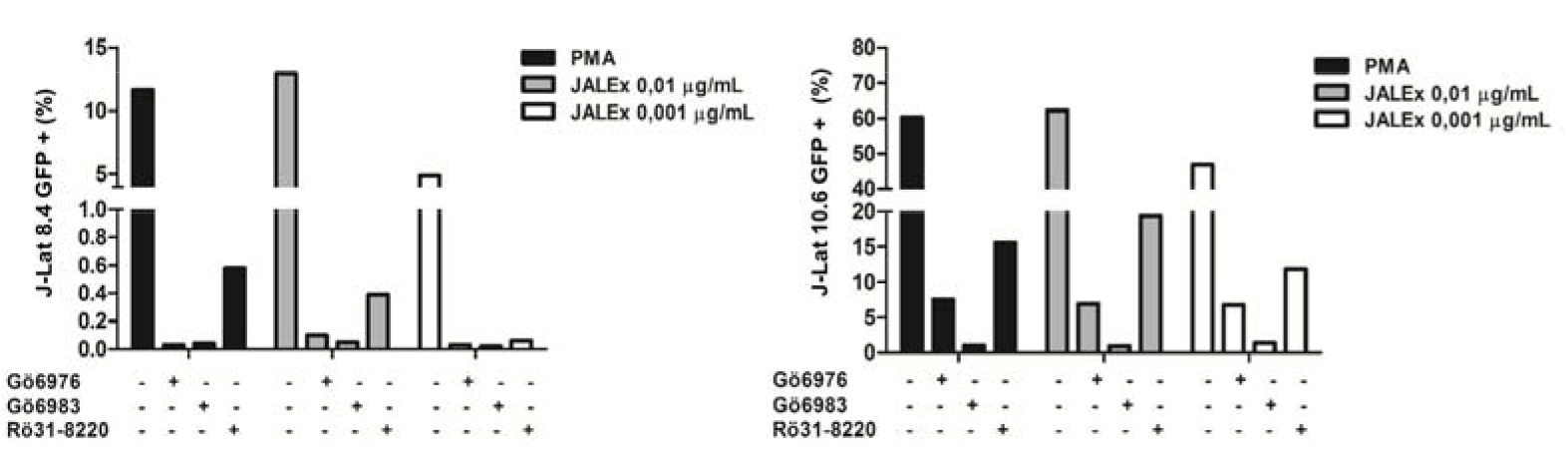
JALEx-mediated HIV-1 reactivation occurs via PKC activation. J-Lat cells 8.4 (left panel) and J-Lat 10.6 (right panel) were pretreated for 24 h with three different PKC inhibitors (G6666, G6363and Ro-31-8220) at the concentration of 1 μM each. The cells were then incubated with different concentrations of JALEx for an additional 24 hours and GFP expression was assessed by flow cytometry. PMA (1 μM) was used as a positive control for activation of PKC-dependent HIV-1 (n=1).

### JALEx induces activation of different isoforms of PKC

PKC family is a citoplasmatic sensor of several signal transduction pathways by phosphorylation of hydroxyl groups in serine/threonine residues of other proteins. Moreover, the catalytic domain present in the structure of PKCs is essential for its auto-phosphorylation and, consequently, its cellular activation and relocation to the inner face of plasma membrane. PKC family comprises different isoforms spanning different regulatory and catalytic domains. The isoforms are divided into three subgroups: the conventional isoforms **α**, **βI**, **βII** and **α** that are Ca^2+^ and DAG dependente for activation; the novel class encompassing the **δ**, **ε**, **η** that require DAG, but not Ca^2+^ for activation and the atypical isoforms **ι** and **λ** that require neither Ca^2+^ nor diacylglycerol for activation. For these reasons, we investigated whether phosphorylation of different PKC isoforms was occurring in response to JALEx treatment. In these experiments, two concentrations of JALEx (0.01 μg/mL and 0.001 μg/mL) were used and PMA was utilized as positive control of PKC phosphorylation. Cells were collected at different time points (10 and 30 minutes; 1, 6 and 24 hours). We observed that the phosphorylation of conventional PKCs, identified by anti-pan-PKC-phosphorylated antibody labeling, occurs quite rapidly, after 10 minutes of treatment with JALEx, in the two concentrations tested (Fig. S2A, upper panels). This activation is long lasting, remaining phosphorylated for up to 24 hours after the treatments (Fig. S2A, upper panels). The same pattern was observed at the same time points for cells treated with PMA (Fig. S2A, upper panels). For the novel PKC isoforms, PKC **θ** exhibited a rapid phosphorylation pattern (10 min) that was not sustained over time (1 to 24h) (Fig. S2A, bottom panels). PKC **δ** showed an early phosphorylation pattern maintained for up to 6 hours with JALEx treatment in both concentrations (0.01 μg/mL and 0.001 μg/mL). However, only JALEx at 0.001 μg/mL induces PKC **δ** phosphorylation after 24 hours (Fig. S2A, bottom panels). Densitometry analyses compared with loading controls (**α**-tubulin) and normalized by DMSO treated cells showed that, in general, JALEx at concentrations of 0.01 μg/mL and 0.001 μg/mL induces the phosphorylation of conventional and novel isoforms of PKC that remains at least up to 6 hours of stimulation (Fig. S2B). However, as suggested by the WB analysis, the phosphorylation of PKC-**δ** is sustained until 24 hours post JALEx treatment. All these results are in agreement with the previous observations that JALEx is a pan activator of PKC isoforms.

activation, PKCs are translocated to the plasma membrane by RACK proteins (membrane-bound receptor for activated protein kinase C proteins) (**46**). To verify if JALEx promotes the translocation of PKC isoforms to the plasma membrane, HeLa cells were transfected with plasmids expressing three PKC isoforms fused to GFP (PKC**α**-GFP, PKC**δ**-GFP and PKC**γ**-GFP). After transfection, cells were treated with 1 and 0.1 μg/mL of JALEx for 10 minutes or 24 hours and the citoplasmatic location of PKC were evidenced by fluorescence confocal microscopy images. We observed a diffuse pattern of PKC distribution in untreated cells (mock). However, PMA or JALEx treatment at both concentrations (1 and 0.1 μg/mL) induced the periplasmatic distribution of all PKC isoforms (**α**, **δ** and **γ**) (Fig. 4A). The same results were obtained with longer treatment scheme (up to 24h) (Fig. 4B). Finally, these results showed that JALEx promotes activation of different classes of PKC through phosphorylation followed by downstream relocation to the plasma membrane.

**Figure 4:**
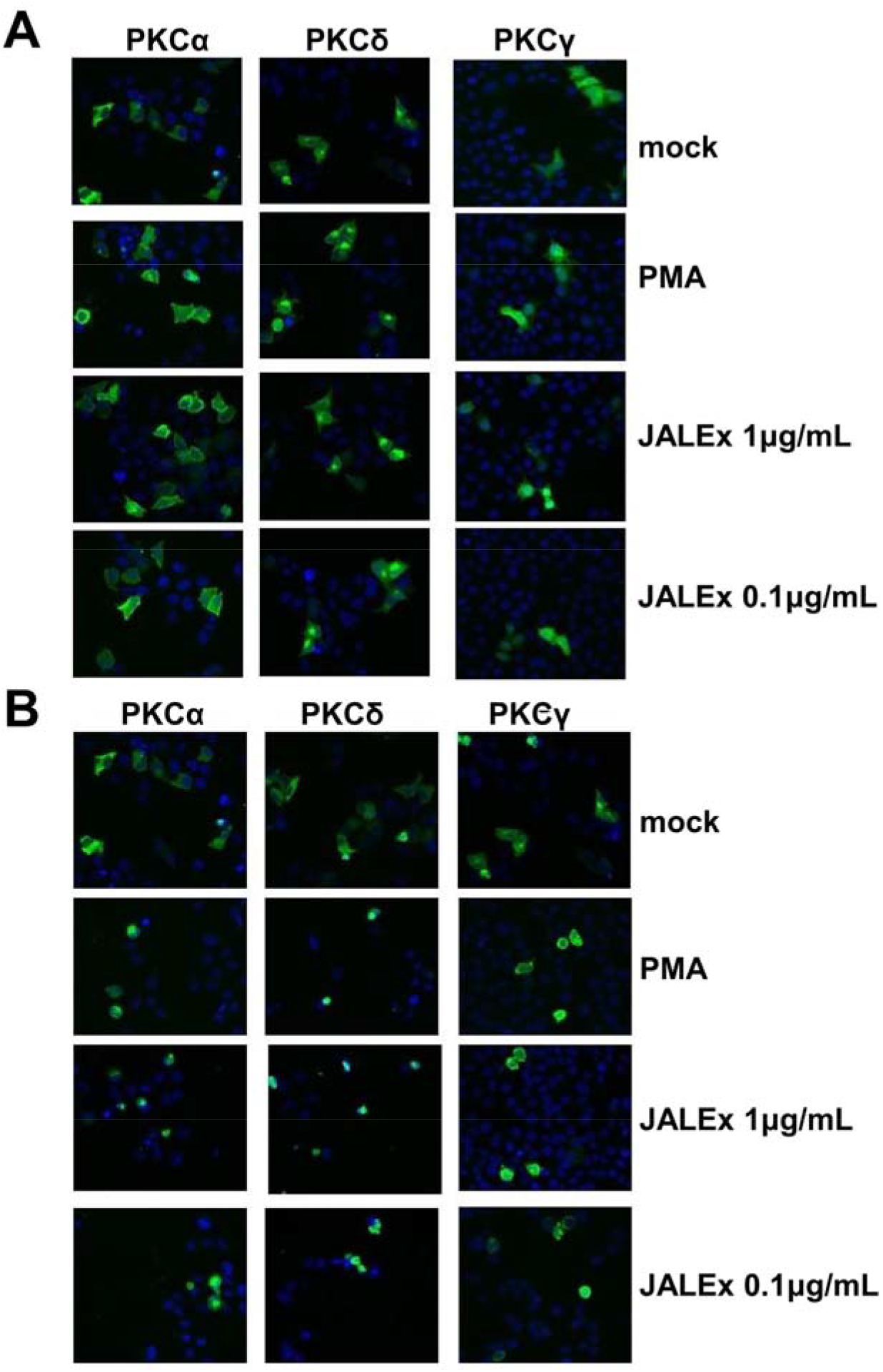
JALEx is responsible for the cellular relocation of PKC isoforms. Cellular location analysis of different PKC (alpha, delta and gamma) isoforms after treatment of HeLa cells with two concentrations of JALEx: 1 μg / mL and 0.1 μg / mL. (A) HeLa cells were transfected with 3 different PKC isoforms for 24 h. After this time the cells were treated for 10 minutes with the two concentrations of JALEx. (B) HeLa cells were transfected with 3 different PKC isoforms for 24 h. After this time the cells were treated for additional 24 hours with the two concentrations of JALEx. As a positive control, PMA (1 μM) was used (n=3).

### JALEx promotes nuclear translocation of NF-κB and interaction with HIV promoter

Phosphorylation of NF-kB inhibitor IκB by PKC leads to its ubiquitination and proteasomal degradation, releasing NF-κB complexes in the cytoplasm to translocate to the nucleus where they bind to HIV promoter (LTR region) and induce virus gene expression (**47–49**). For this reason, we evaluated the effect of JALEx treatment on nuclear translocation of NF-κB by means of two strategies. In the first one, we analyzed the occurrence of nuclear translocation of NF-κB after treatment of HeLa cells with 1 and 0.1 μg/mL of JALEx for 6 or 24 hours, through immunofluorescence and confocal fluorescence microscopy using anti-p55 antibody. We observed no significant translocation of NF-κB to the nucleus of HeLa cells 6 hours post JALEx treatment with the same cytoplasmic distribution as observed in untreated cells (mock - Fig. 5A). However, we confirmed the NF-kB translocation to the nucleus after 24 hours of JALEx treatment, once NF-kB staining was located in the nuclei (Fig. 5B). To better quantify the translocation of NF-κB, we quantified the mean values of fluorescence intensity between the nucleus and the cytoplasm of 100 cells, in each experimental condition. We observed an increasing ratio of NF-kB fluorescence into the nucleus 24 hours after JALEx treatment, and the same was observed for the positive control (PMA - Fig. 5C). JALEx treatment at the concentration of 1 μg/mL showed the highest rates of NF-kB internalization (Fig. 5C).

**Figure 5:**
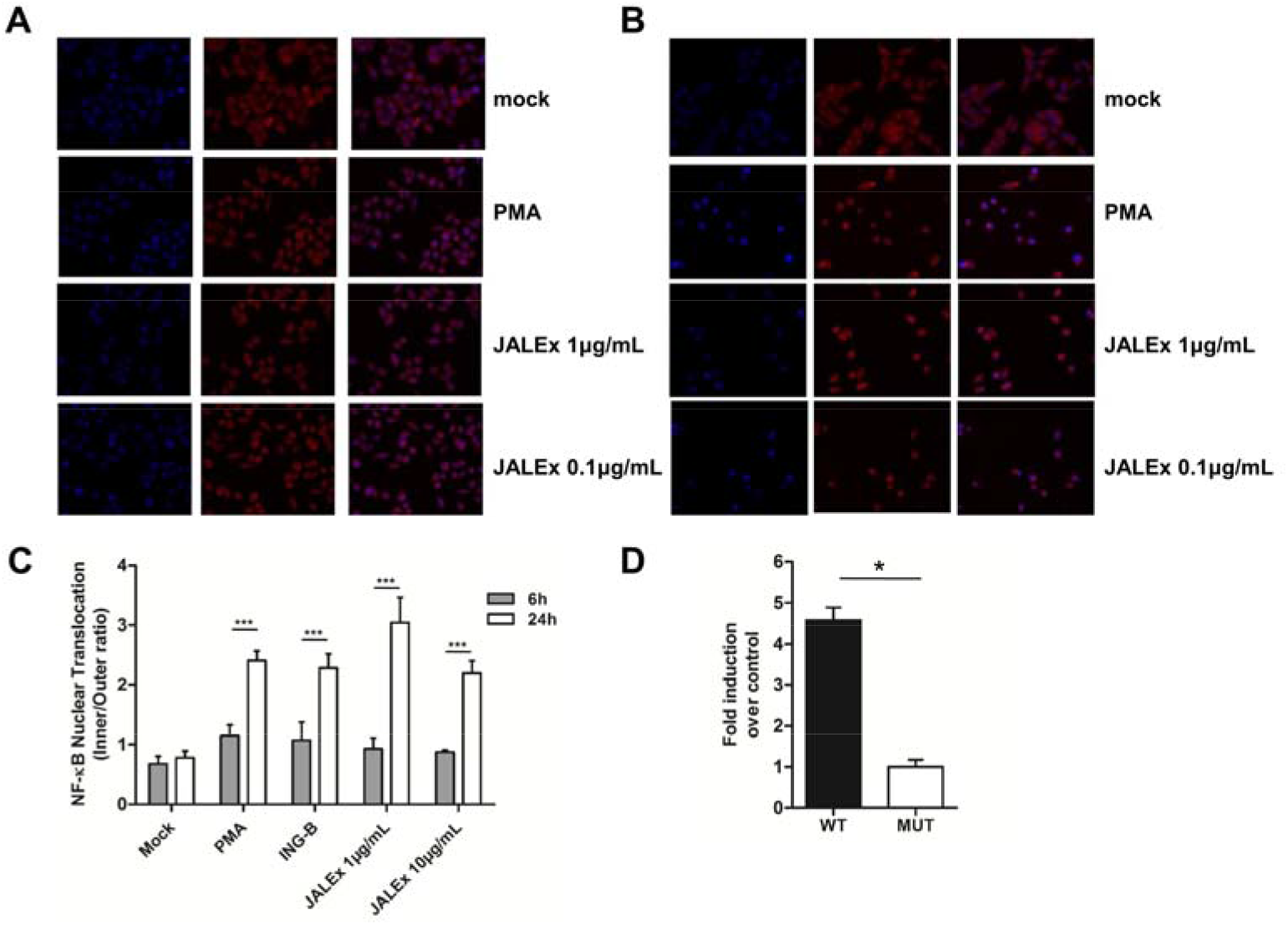
JALEx-mediated transcriptional activation is dependent on NF-κB activation. HeLa cells were treated with two concentrations of the **JALEx** (1 and 0.1 μg/mL) for 6 hours (A) and 24 hours (B). After these time intervals, the cells were subjected to immunofluorescence for labeling of NF-κB, in red and cell nucleus, in blue. As a positive control, PMA (1 μM) was used. (C) Graph of the quantification of cells showing NF-κB translocation. An average value of the fluorescence intensity between the nucleus and the cytoplasm of 100 HeLa cells was calculated under all conditions, at the time intervals of 6 and 24 h after the treatments (n=3; Two-way ANOVA and (***) indicate p<0,0001). (D) Jurkat cells were transfected with pBlue30LTR-Luc (NF-κB WT) and the mutant construct, pBlue30LTR NF-κB MUT-Luc (NF-κB MUT), absent from the NF-κB binding site, preventing its interaction with NFκB. After transfections, the cells were treated with 1 μg / mL of the **JALEx** for 24 hours and luciferase expression was measured after lysis of the cells. The fold-change of the luciferase expression between the wild-type plasmid and the mutant is shown in the graph (n=4, Mann-Whitney test and (*) indicate p<0,05). The bars in Figure 5 C and D represents 1 SD around the mean.

We also evaluated the recruitment of NF-κB to the HIV promoter by JALEx treatment. For that, we transfected Jurkat T CD4^+^ cells with LTR-luciferase reporter plasmids containing either intact NF-κB binding sites (NF-kB WT) or harboring mutations that prevent NF-κB interaction (**17, 50, 51**). Transfected cells were treated with 1 μg/mL of JALEx and luciferase activity was assessed 24 hours later. Our results showed that JALEx treatment induced a 5-fold increase in luciferase expression in cells transfected with pNF-kB WT. However, no luciferase activity was observed in cells transfected with pNF-kB MUT (Fig. 5D). Together, our results demonstrated that JALEx induces HIV transcription through NF-kB release of its inhibitors, followed by nuclear translocation and binding to HIV promoter.

### JALEx modulates surface markers expression and cytokines release in primary human CD4^+^ lymphocytes

To analyze whether the effects of treatment with JALEx would have an impact on primary human cells, we performed assays on CD4^+^ T cells isolated from PBMCs from 4 healthy donors. Isolated TCD4^+^ cells were treated with PMA (1 μM) or JALEx 10 μg/mL or 1 μg/mL for 24 hours. We evaluated the expression of HIV receptor and co-receptors (CD4, CCR5 and CXCR4) and activation markers (CD25, CD38, HLA-DR and CD69) in response to JALEx treatment. We observed that JALEx treatment (1 μg/mL) downregulates the surface receptors CD4 and CXCR4 expression levels without any alteration in CCR5 expression levels (Fig. 6A). The same pattern was observed for cells treated with PMA (Fig. 6A). Indeed, JALEx treatment also induced the expression of surface activation markers CD25, CD38 and HLA-DR when compared to non-treated cells, and with PMA control (Fig. 6B). Figure 6C shows that CD69 activation marker expression is 75-fold higher relative to control cells in both treatments. These results suggest that JALEx not only reactivate latent HIV, but also could prevent new cycles of virus infection down modulating the virus receptors and co-receptors.

**Figure 6:**
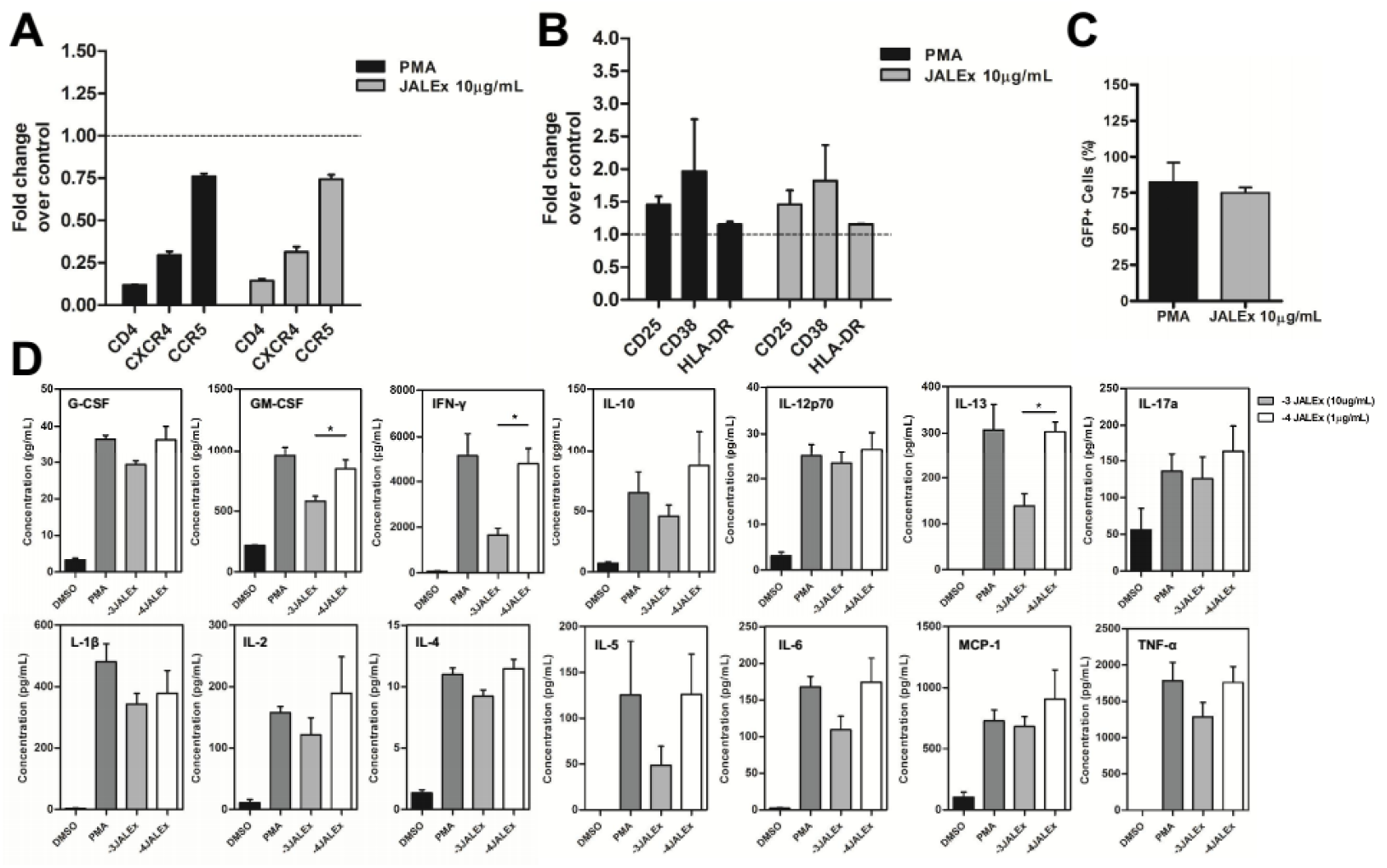
JALEx modulates the expression of cell surface receptors, activation markers and cytokine secretion in primary CD4 + T cells from HIV-individuals. TCD4 + cells were isolated from PBMCs from 4 healthy donors and treated for 24 hours with JALEx. After this period, the expression of some surface molecules was evaluated. (A) Expression of the CD4 receptor and CCR5 and CXCR4 co-receptors in CD4 + T cells isolated after treatment with JALEx (n=4). (B) Expression of CD25, CD38 and HLA-DR activation markers in CD4 + T cells isolated after treatment with JALEx (n=4). (C) Expression of the CD69 activation marker on CD4 + T cells isolated after treatment with janauba extract. The black bar represents the positive control (PMA 1 μM) and the gray bar represents the treatment with 10 μg / mL of JALEx (n=4). (D) CD4+ T cells were treated with 1 μM PMA, 10 μg/mL (−3) or 1 μg/mL (−4) of JALEx for 24 hours. After, supernatants from the CD4+ T cells were collected and cytokine analysis were performed by Bio-plex Pro Human Cytokines 17-plex platform (n=5, Kruskal-Wallis test and (*) indicate p<0.05). The bars in the figure represents 1 SD around the mean.

In the light of the previous results of JALEx inducing PBMC activation, we further analyzed cytokines release directly from the supernatant of human CD4^+^ T cells. This analysis was performed using a multiplex immunoassay to quantify the production of 17 pro-inflammatory cytokines. JALEx treatment induced the production of the following cytokines relative to DMSO treated cells: IL-1**β**, IL-2, IL-4, IL-5, IL-6, IL-10, IL-12 (p70), IL-13, IL-17, G-CSF, GM-CSF, MCP-1/MCAF, IFN**γ** and TNF-**α** (Fig. 6D). Overall, the highest concentration of JALEx (10 μg/mL) induced the same levels of cytokines as did PMA positive control. IFN-**γ** and TNF-**α** were highly induced under JALEx treatment, 5,000 and 1,800 pg/mL, respectively (Fig. 6D). Those results showed that JALEx activate T CD4^+^ cells in order to secrete important pro-inflammatory cytokines involved in macrophage recruitment (MCP-1) and T CD8 response (INF-**γ**) to block HIV replication. MIP-**β**, IL-7 e IL-8 levels were undetectable.

### JALEx increases viral load in PBMC from HIV-1 positive patients and induces Th1 and Th17 immune response

Progression for AIDS in HIV-1 positive patients is mainly characterized by CD4^+^ T cell depletion specifically Th1 and Th17 cells, which impact negatively on immune protection against different opportunistic pathogens. We first analyzed the virus activation on PBMC isolated from HIV-1 positive patients under ART treatment. We found that JALEx at 1 μg/mL increases HIV viral load up to 1.5 log compared with non treated PBMC cells (Fig. 7A). This data was significant in all PBMC generated from the 15 HIV positive patients. This increase in HIV viral load was followed by T CD4^+^ cells depletion (up to 82%) with no correspondence in T CD8^+^ cells (Fig. 7B). In order to investigate the immune response in those patients we evaluate cytokine production in T CD4 and T CD8 positive cells. JALEx treatment (0.1 μg/mL) increased the frequency of CD4^+^ and CD8^+^ T cells secreting IL-17, IFN-_γ_ and IL-21 cytokines. This induction was higher compared to PMA positive controls (Fig. 7C). Moreover, JALEx (0.1 μg/mL) enhanced not only the percentage of single IL-21-secreting CD4^+^ and CD8^+^ T cells, but also Th1 and Th17-like subsets positives for IL-21 (Fig. 7D). In contrast, no difference was observed regarding either IFN-_γ_ or IL-17-secreting CD4^+^ and CD8^+^ T cells from PBMC cultures of HIV-1 positive patients following JALEx treatment (0.1 μg/mL, Fig. 7D). These results suggested that JALEx treatment could reactivate HIV from reservoirs in patients with ART suppressive therapy and induces CD4 and CD8 positive cells secreting protective levels of IL-21.

**Figure 7.**
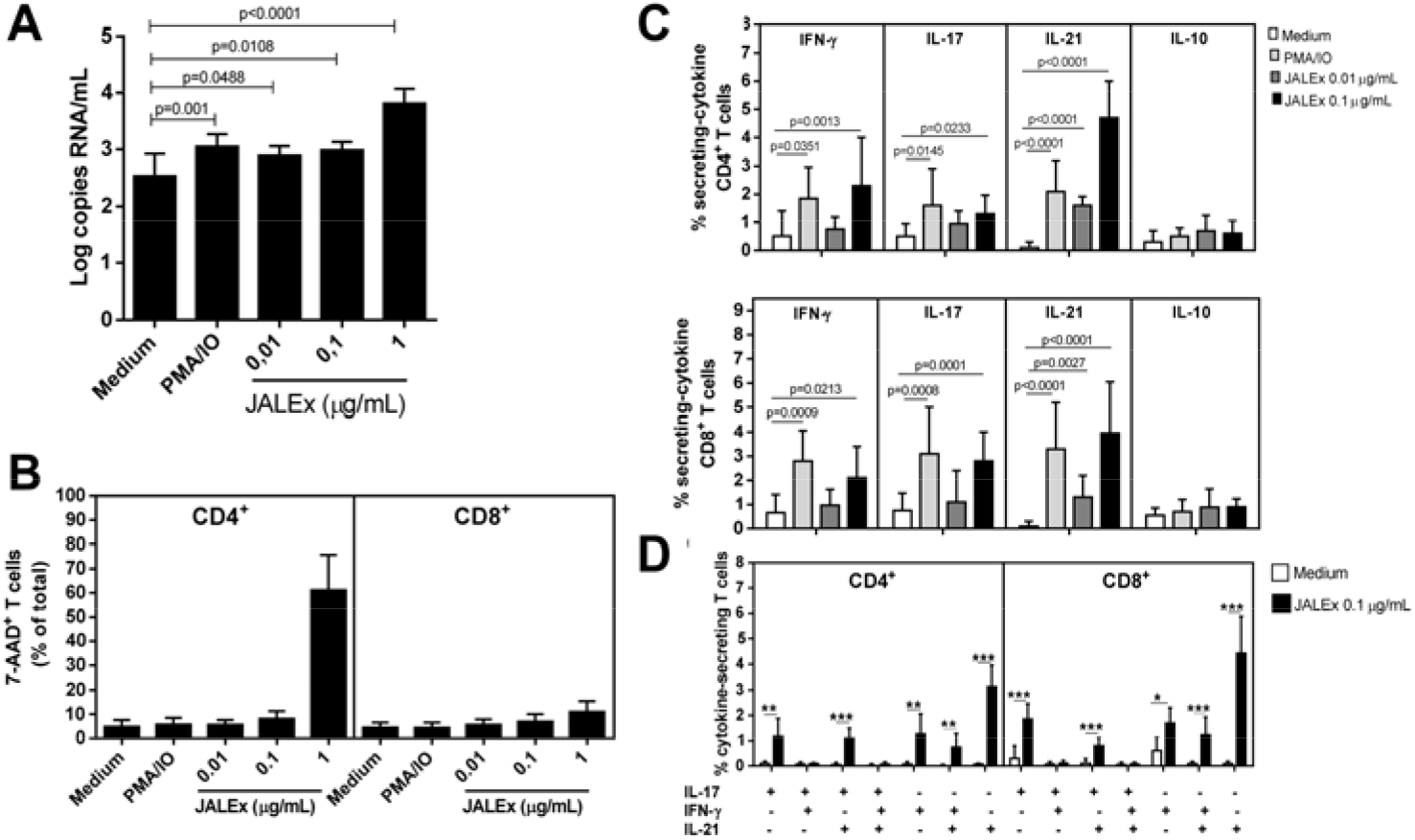
Effect of JALEx on T cell viability, HIV-1 replication in PBMC cultures and cytokine secretion in T cells from AIDS patients. Different concentrations of JALEx (0.01, 0.1 and 1 μg/mL) were added to PBMC cultures (1 × 10^6^/mL) obtained from 15 AIDS patients, and 24h after, the (A) the number of HIV-1 into supernatants was determined through RT-qPCR, (B) CD4^+^ and CD8^+^ T cells viability was analyzed by FACS after 7-AAD staining. AIDS-derived PBMC cultures (1 × 10^6^/mL, n=15) were maintained for 24h in the presence of medium alone or with different JALEx ( 0.01 and 0.1 μg/mL). As positive control, some well receiving PMA plus ionomycine (PMA/IO). The percentage of CD4+ and CD8+ T cells producers of IFN-**γ**, IL-17, IL-21 and IL-10 was determined by cytometry (C). The mean values were compared and the p values are shown in the figures. In (D), the proportion of different (CD4+ and CD8+) T cell subsets of secreting different combinations of IL-17, IFN-**γ** and IL-21 in response to JALEx (0.1 μg/mL) was also evaluated by cytometry. In the figure, (*), (**), and (***) indicate p<0.05, p<0.001 and p<0.0001, respectively. The bars in the figure represents 1 SD around the mean.

### JALEx reactivation effects in latently SIV-infected macaque cells

Finally, we explored the capacity of JALEx to reactivate latent viruses in CD4^+^ lymphocytes from SIVmac239-infected *cynomolgus macaques*. We purified resting memory CD4^+^ T cells from a single viremic macaque and prepared serial dilution replicates of these cells to seed into quantitative viral outgrowth assays (QVOA). Cells in these assays were stimulated in triplicate at each cell dilution with JALEx or other compounds known to reactivate latent HIV/SIV through PKC pathway (ingenol and concanavalin A). In order to compare the ability of these compounds to reactivate latent SIV, we measured the frequency of infected cells (quantified as infectious units per million cells; IUPM) across treatments. A higher frequency of infected cells after stimulation with a given compound suggests its efficiency to reverse latency. The highest frequency of infected cells was observed in cultures treated with JALEx, although this frequency was not significantly different from those observed in other cultures (Table S1). However, JALEx was the only compound capable of reactivating SIV in all replicates tested showing its superiority compared with others PKC agonists, such as Ingenol and Concanavalin A.

## DISCUSSION

In this work we addressed the ability of an ethanolic extract obtained from *Euphorbia umbellata* latex (JALEx) to reactivate HIV/SIV latency in several *in vitro* experiments. The traditional medical uses of this Euphorbiaceae are well known in South America and this plant is vastly used diluting a small volume of crude latex in drinkable water and consuming this solution regularly (**24**). JALEx contains a series of bioactive phorbol esters (see Supplemental Figure 1). These class of molecules such as PMA are known PKC agonist implicated in lymphocyte activation as well HIV reactivation in well conducted *ex vivo* studies and these activities requires both PKC**α** and PKC**θ** isoforms activation (**52**). Modifications in phorbol ring can lead molecules able to induce carcinogenesis and others such as a deoxy in ring position 4 (4-deoxyphorbol) can lead to molecules unable to induce carcinogenesis but retaining its PKC activation capacity (**53**). The presence of this modified phorbol ester was identified in our JALEx in our chemical analysis and ina previous study done with *Euphorbia umbellata*latex (**45**). Therefore, our JALEx is not likely to induce cell carcinogenesis retaining the capacity to reactivate HIV/SIV latency. Many LRA act activating PKC and several compounds such as prostatin (**15**), bryostatin (**16**), ingenol B (**17**), and phorbols such as PMA (**18**). Fit in this class of molecules Prostratin and bryostratin, are strong PKC agonist and they are very toxic when injected intravenously. In fact, bryostatin was used in a phase II study with 17 patients with progressive non-Hodgkin’s lymphoma of indolent type (NHL), previously treated with chemotherapy. These patients received a median of 6 intravenous infusions of 25□μg/m^2^ bryostatin 1 given once weekly over 24 h and the principal toxicities were myalgia and phlebitis (**54**). Ingenol B a semisynthetic compound isolated from *Euphorbia tirucalli* was able to reactivate HIV *ex vivo* with great potency using the PKC/NF-kB pathway inducing increased cellular levels of CycT1 and CDK9 in human CD4+lymphocytes in ex vivo experiments (**17**).

In our study we used JALEx to explore its capability to reactivate HIV latency taking the advantage of the use of this plant in traditional medicine without serious side effects. First we used JALEx in J-Lat 8.4 and 10.6 cells and we could show the potency of this extract reactivates latent HIV transcription present in J-Lat 8.4 and 10.6 cells in low concentrations (0.01 μg / mL).We also demonstrate that JALEx act through PKC by inhibiting J-Lat activation with three different PKC inhibitors (G6666, G6983 and Ro-31-8220). In our assays Gö6983 was more powerful inhibitor totally blocking HIV reactivation in both J-Lat cell lines. Go 6983 is a fast pan-PKC inhibitor against PKC**α**, PKC**β**, PKC**δ** and PKC**δ** with IC_50_ in low nM range then JALEx could be reactivating latent HIV through the activation of different PKC isoforms. Additionally, we showed that JALEx activates the PKC pathway by promoting NF-κB internalization into the nucleus and increasing HIV transcription, which is dependent on NF-κB binding to LTR promoter. Using image and western blot analysis we showed that JALEx was able to activate classical and novel PKC isoforms such as: the conventional isoforms **α**, **βI**, **βII** and **γ** as well the new ones encompassing the **δ**, **ε**, **η** and **θ** isoforms in *in vitro* experiments using JURKAT cell lines. JALEx could induce **θ** isoform, in high levels. The transcription factors NF-κB and AP-1 are the primary physiological targets of PKC**θ**, and efficient activation of these transcription factors by PKCθ requires integration of TCR and CD28 costimulatory signals. PKCΘ cooperates with the protein Ser/Thr phosphatase, calcineurin, in transducing signals leading to activation of JNK, NFAT, and the *IL-2* gene. PKC**θ** also promotes T cell cycle progression (**57**).

We also evaluate the capacity of JALEx to induce cytokines production in human PBMC cells *in vitro*. This pattern of alteration in the production of cytokines from treatment with reactivating compounds is quite common in the case of the shock and kills strategy, which aims to activate the cells and promote the exit of HIV latency that is integrated into the cells T. This phenomenon is known as “cytokine storm” and needs to be further studied in animal models to define a dose that does not induce this toxic reaction. Although JALEx induced a conjunction of pro inflammatory cytokines we could observed a noticeable level of induction of IL-21 by different CD4+ and CD8+T lymphocyte subsets from PBMC cultures from Aids patients, a noble cytokine enrolled in the HIV virus load control in HIV+ individuals (**55, 58, 59**). For this reason, the search for new, more selective compounds that target only a few PKC isoforms or that have their mechanism of action in other stages of the activation pathways to reduce these risks is necessary. Of note, leaf and latex ethanolic extract of E. umbellata latex had low toxicity in adults Rat when dose orally and acute doses of 2000mg/kg given orally was well tolerated (**56**).

In our experiments JALEx was efficient in reactivating HIV production directly in resting T CD4+ cells in HIV+ individuals in ex vivo assays measuring viral RNA in cell supernatant as well primary non-human primate T CD4+ cells counterpart in SIV251 latency model using a QVOA assay in low concentrations. The anti-HIV action of JALEx could occur by three synergistic effects: a)reactivation of HIV from latency and its destruction by ART; b) blocking infection of new cells through the down-modulation of CD4 on the lymphocyte surface; and c) by inducing the secretion of antiviral cytokines such as IL21. All these *in vivo* properties of JALEx in non-human primate cell models added to the molecular mechanisms of reactivation presented here as well the traditional use of *E. umbellata* latex, strongly recommends JALEx or other *E. umbellata* latex preparations as a great candidate for future clinical trials for HIV eradication

## Acknowledgments

We thanks Dr. Thomas Friedrich from Virology Services Unit at the Wisconsin National Primate Research Center, Wisconsin, USA for QVOA assay using SIVmac239 in NHP CD4^+^ Lymphocytes. We are also grateful to Mr. Wilson Camargo Barros Filho and Mr. Luis Antônio Nogueira for providing *E. umbellata´s* latex used in this work. This work was sponsored by CNPq, FAPERJ, CAPES and FINEP.

## SUPPLEMENTAL MATERIAL

**Figure S1:**
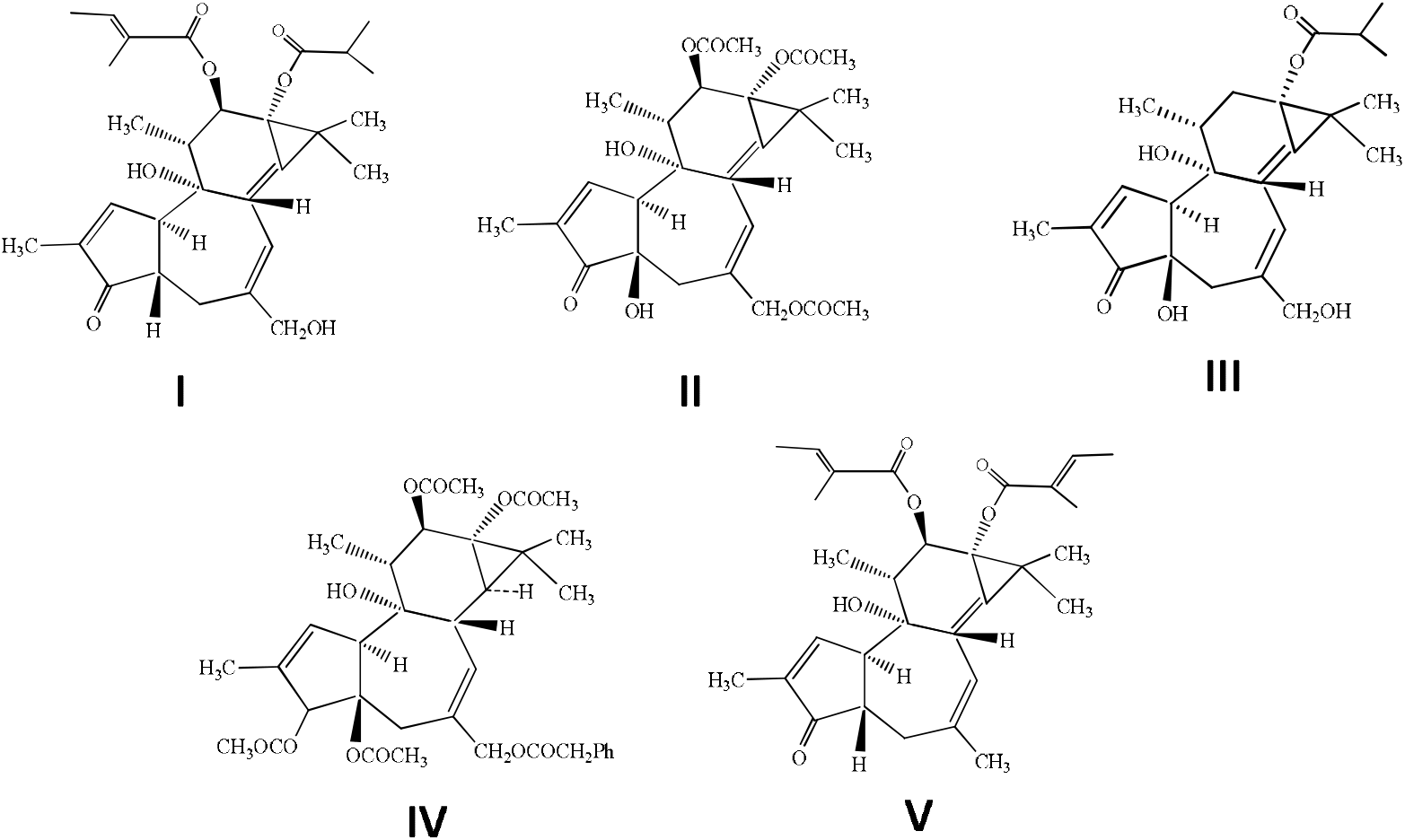
Phorbol esters isolated from *Euphorbia umbellata*. I = 12-*O*-Tigloyl-4-deoxyphorbol-13-isobutyrate; II = Phorbol-12,13,20-triacetate; III = 12-Deoxyphorbol-13-(12-methylpropionate); IV = 3,4,12,13-Tetraacetylphorbol-20-phenylacetate (synagrantol A); V = Deoxyphorbol-12,13-ditiglate (synagrantol B).

**Figure S2:**
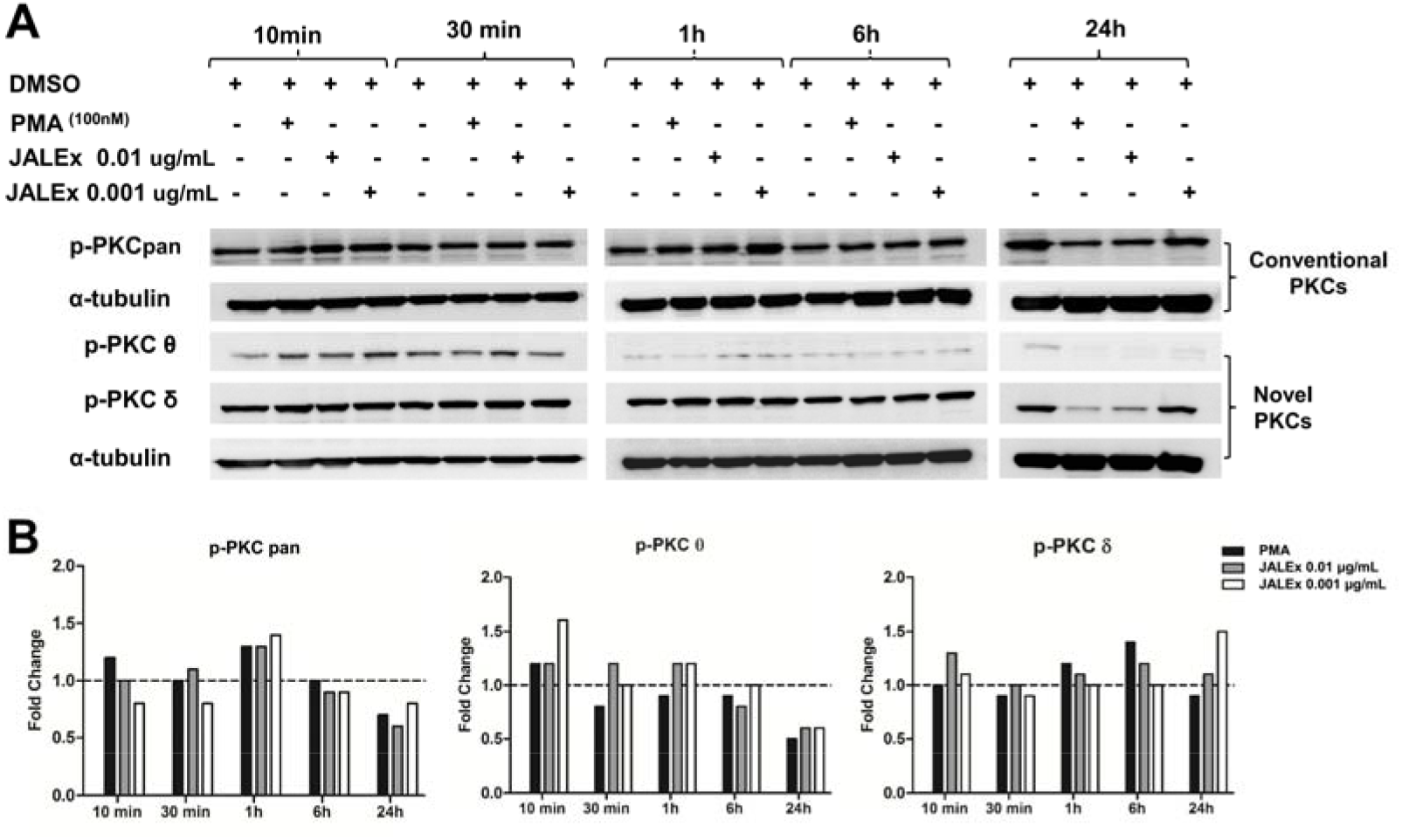
Panel of phosphorylation of different PKC isoforms after treatment with JALEx. (A) Jurkat cells were treated with two different concentrations of JALEx (0.01 μg/mL and 0.001 μg/mL), PMA (1 μM) as positive control, at different time intervals (10, 30 minutes, 1, 6 and 24 hours). Then the cells were lysed for western blotting with phosphorylated anti-PKC antibodies (pan, **δ**, **θ**) and anti-tubulin as the loading control. (A) The intensity of the western blotting bands corresponding to the Jurkat cells in (B) that were quantified by densitometry with the aid of the Image J program. Dashed lines correspond to the band intensity of MOCK in these experiments (n=1).

**Table S1.** QVOA results using macaques CD4+ T Lymphocytes extracted from SIV 251 infected Macaques treated with HAART therapy.

| Cells/well<sup>a</sup> | No mitogen | Con A | Ingenol B | **JALEx** |
| --- | --- | --- | --- | --- |
| 1.25e5 | + / - / -<sup>b</sup> | - / - / - | + / - / + | + / + / + |
| 6.25e4 | - / - / + | - / + / + | + / + / - | + / + / + |
| 3.1e4 | + / - / + | - / - / + | - / + / + | + / + / + |
| Maximum likelihood (IUPM) | 7.735675 | 5.229369 | 15.525127 | **34.986678** |
| 95% CI | {2.866090, 20.878852} | {1.680200, 16.275622} | {6.619641, 36.411279} | {12.619641, 66.411279} |

<sup>a</sup>- Number of CD4+ T Lymphocytes extracted from SIV 251 infected Macaques treated with HAART therapy inoculated in each well. <sup>b</sup>- Number of well with p27 antigen positive. Con A:

